# Efficient Analysis of Annotation Colocalization Accounting for Genomic Contexts

**DOI:** 10.1101/2023.11.22.568259

**Authors:** Askar Gafurov, Tomáš Vinař, Paul Medvedev, Broňa Brejová

## Abstract

An annotation is a set of genomic intervals sharing a particular function or property. Examples include genes or their exons, evolutionarily conserved elements, and regions with a particular epigenetic state. A common task is to compare two annotations to determine if one is enriched or depleted in the regions covered by the other. We study the problem of assigning statistical significance to such a comparison based on a null model representing two random unrelated annotations. To incorporate more background information into such analyses, we propose a new null model based on a Markov chain which differentiates among several genomic contexts. These contexts can capture various confounding factors, such as GC content or assembly gaps. We then develop a new algorithm for estimating p-values by computing the exact expectation and variance of the test statistics and then estimating the p-value using a normal approximation. Compared to the previous algorithm by Gafurov et al., the new algorithm provides three advances: (1) the running time is improved from quadratic to linear or quasi-linear, (2) the algorithm can handle two different test statistics, and (3) the algorithm can handle both simple and context-dependent Markov chain null models.

We demonstrate the efficiency and accuracy of our algorithm on synthetic and real data sets, including the recent human telomere-to-telomere assembly. In particular, our algorithm computed p-values for 450 pairs of human genome annotations using 24 threads in under three hours. Moreover, the use of genomic contexts to correct for GC bias resulted in the reversal of some previously published findings.

**Availability:** The software is freely available at https://github.com/fmfi-compbio/mcdp2 under the MIT licence. All data for reproducibility are available at https://github.com/fmfi-compbio/mcdp2-reproducibility

## 1 Introduction

Recent years have brought rapid growth in the number of different assays that can extract genome-scale functional information. This has led to growing collections of genome annotations; for example in the UCSC Genome browser, the GRCh38 human genome assembly features thousands of different annotation tracks. In this work, we provide new models and algorithms for annotation colocalization analysis, where the goal is to determine if one input annotation is significantly colocated with regions covered by another annotation. Such analyses may hint at possible connections between biological processes governing individual annotations. For example, sites marked by histone modification H3K4me3 are colocated with promoter regions, and H3K4me3 indeed plays a role in gene transcription regulation (Guenther et al., 2007).

Mathematically, we view a genome annotation as a set of non-overlapping chromosomal intervals. Given two annotations, query *Q* and reference *R*, we consider two widely-used colocalization statistics. The *overlap statistic* is the number of intervals in *R* that intersect with at least one interval in *Q*. The *shared bases statistic* is the number of genomic positions covered by both *R* and *Q*. However, even randomly generated annotations will overlap by chance. In order to ascertain statistical significance of the observed statistic, its p-value needs to be computed under a suitable null hypothesis. Until very recently, all the methods (Dozmorov et al., 2016; Gafurov et al., 2022; Gel et al., 2016; Heger et al., 2013; Sandve et al., 2010; Sarmashghi and Bafna, 2019; Sheffield and Bock, 2016) were limited by having a null hypothesis that either does not properly model the data or its p-value computation does not scale to annotations of human-sized genomes.

Recently, Gafurov et al. (2022) proposed an alternative null hypothesis in which the annotation is produced by a two-state Markov chain. The algorithm, called MCDP, was a substantial improvement in time and memory over previous approaches. However, it is quadratic in the number of reference intervals and still takes several hours for a human exon reference annotation. It thus remains time-prohibitive to compare many pairs of annotations against each other.

Another limitation of MCDP as well as other approaches is that two unrelated annotations may appear to be colocalized because they are each colocalized with another genomic feature (Kanduri et al., 2019). For example, two annotations may appear colocalized simply due to their prevalence in regions with high GC content, even though they are not related. More generally, different regions of the genome can be thought of as providing different background to the null model, and we think of these various backgrounds as partitioning the genome into *contexts*. Accounting for contexts in calculating p-values is important to limit false associations, yet this capability is limited or absent in existing tools.

In this paper, we propose a model and an algorithm to overcome these scalability and accuracy barriers. Our first contribution is a new algorithm MCDP2 for estimating p-values, which is linear in the number of reference intervals. To demonstrate the scalability of our algorithm, we considered 10 reference annotations, corresponding to different types of repeats in the human genome, and 45 query annotations, corresponding to epigenetic modifications in different cell lines. MCDP2 computed p-values for all 450 pairs using 24 threads in under 2 hours for the number of overlaps and 3 hours for the number of shared bases.

Our second contribution expands the modeling capability of the Markov chain null hypothesis so that it takes into account genomic context and thus captures various confounding factors influencing annotation colocalization. Unlike previous approaches (Heger et al., 2013), our model is able to handle annotation intervals that span class boundaries. We demonstrate the importance of modeling the genomic context by re-analyzing colocalization of copy number deletions with various gene classes (Zarrei et al., 2015) and find that adding a genome context in fact reverses some of the previous conclusions. In one striking example, the set of all exons appears enriched for overlap with copy number losses but enrichment turns into depletion after taking into account gaps and GC content. We also compare the colocalization of epigenetic marks with subtelomeric repeats on the new human telomere-telomere assembly (Gershman et al., 2022), using a genome context to compare enrichment between two classes of repeats.

### Related work

Several null hypotheses for colocalization statistics have been considered prior to the Markov chain model (Gafurov et al., 2022). Some lend themselves to fast and simple statistical tests (e.g. Fisher’s exact test) but do not capture relevant properties of the data. For example, one can assume that all positions in the query annotation are chosen uniformly at random (Dozmorov et al., 2016; Sheffield and Bock, 2016). However, this does not capture either the integrity of intervals or their length distribution. A more faithful option is the permutational null hypothesis (also called gold null hypothesis (Gafurov et al., 2022)), which reshuffles the query intervals while maintaining their lengths (Gel et al., 2016; Heger et al., 2013; Sandve et al., 2010). Computing the exact p-values for the overlap statistic in this model is NP-hard (Gafurov et al., 2022), and the only known efficient algorithms are either inaccurate or impractical for human-sized annotations (Sarmashghi and Bafna, 2019). Sampling approaches can be used, but their accuracy is directly proportional to the number of samples, making it difficult to estimate small p-values. With these limitations, it was impossible to compute small p-values for human-sized genomes while having a null hypothesis that is faithful to the data.

Accounting for genomic contexts has also been considered but most previous approaches (Dozmorov et al., 2016; Gel et al., 2016; Quinlan and Hall, 2010; Sheffield and Bock, 2016) are only able to account for contexts which are completely inadmissible to annotations (e.g. assembly gaps, which are unassembled regions of the genome). A notable exception is GAT (Heger et al., 2013), which splits a genome into multiple contexts and analyzes colocalization in each context independently. However, this approach does not satisfactorily handle intervals that span context boundaries, which become prevalent when the contexts are short.

## 2 Methods

In this section, we define our context-aware Markov chain null model 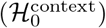 and describe our algorithm for efficient estimation of p-values under this model. We first present our results on a single chromosome; an extension to multiple chromosomes is discussed in Section 2.5.

We will denote the chromosome length as *L* and use 0-based coordinates. An *annotation* is a set of intervals contained in [0, *L*) so that each two intervals are disjoint and separated by at least one base. By |Q| we denote the number of intervals in annotation *Q*. We will represent an annotation either as a list of half-open intervals ordered from left to right *Q* = ([*b*_1_, *e*_1_), …, [*b*_|*Q*|_, *e*_|*Q*|_)), or as a binary sequence *Q* = (*Q*_0_, *Q*_1_, …, *Q*_*L−*1_), where *Q*_*i*_ is 1 if position *i* is covered by one of the intervals and 0 otherwise.

Let *R* and *Q* be two annotations, denoted as the reference and the query, respectively. A *test statistic* is a function that measures the extent to which *R* and *Q* are colocalized. We will consider two concrete test statistics in this paper. One is the number of overlaps *K*(*R, Q*), which is defined as the number of intervals in R that overlap some interval in *Q*. The other is the number of shared bases *B*(*R, Q*), which is defined as the number of bases in the genome covered by both *R* and *Q*. Let *C* be the distribution of the query annotation under the null hypothesis of the query being generated independently of the reference annotation. Given some test statistic *A*(*R, Q*), we are interested to compute the p-value measuring the statistical significance of enrichment of *Q* with respect to *R*, that is, probability Pr_*Q′∼C*_ [*A*(*R, Q*′) *≥ A*(*R, Q*)].

Our algorithm is based on the observation that the distribution of the test statistics is in most realistic scenarios well approximated by the normal distribution (see Sections 3 and 4). Therefore, instead of computing the full probability mass function (PMF), we compute only its exact mean and variance and use them as the parameters of the normal distribution. This means that we calculate the p-value by first computing the *Z-score*, which is the number of standard deviations that *A*(*R, Q*) is above the expected value, under the null. Formally,

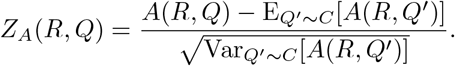

Under the assumption that the statistic is normally distributed, the desired p-value is then simply 1 *−* Φ(*Z*_*A*_(*R, Q*)), where Φ is the cumulative distribution function of the standard normal distribution. Analogously, the p-value for the statistical significance of depletion, defined as Pr_*Q*′∼*C*_ [*A*(*R, Q*^*′*^) *≤ A*(*R, Q*)], is computed as Φ(*Z*_*A*_(*R, Q*)).

In Section 2.1, we describe our context-aware Markov chain model for generating random annotations and then use it to formally define the 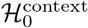 null model in Section 2.2. In Sections 2.3 and 2.4, we describe our algorithm for computing the mean and variance of the overlap and shared bases test statistics, and we also extend it to a more general class of test statistics. Finally, we show how our model naturally extends to multiple chromosomes (Section 2.5).

### 2.1 A generative model

An annotation of a chromosome of length *L* can be generated by running a two-state Markov chain for *L* steps. The state at step *i* indicates whether the annotation includes position *i* on the chromosome. The lengths of the generated intervals and of the gaps between them are known to be geometrically distributed in this model, and the transition probabilities of the Markov chain dictate the expected values of these two distributions (Koller and Friedman, 2009). The Markov chain generative model makes many properties easy to derive and fast to compute (Gafurov et al., 2022), and so we build upon it in this work.

We want to use such a generative model to test if a given query annotation *Q* behaves as if it was “randomly shuffled” on the chromosome. To this end, we set the parameters of the Markov chain so that the expected interval lengths and gaps between them match what is observed in the query *Q*. However, this does not allow to incorporate background knowledge of the chromosome; i.e., some regions of the genome may be *a priori* more likely to contain an interval.

We therefore introduce the notion of genome contexts. Given a finite set of class labels Λ, a *genome context* is a mapping *ϕ* : {0, …, *L −* 1} *−→* Λ of each position on the genome onto a class label (e.g. Λ = {gap, non-gap}). This mapping partitions the genome into several segments with the same class assigned. We will refer to the positions where the class differs from the class label at the previous position as to *class boundaries*. We assume throughout the paper that a context is represented as a sequence of class boundary positions with the corresponding class labels, sorted in an increasing order by positions.

Our generative model allows each context class to have its own Markov chain, i.e. its own distribution of interval lengths and gaps. An annotation is then generated by iterating over the genome positions from left to right, and at each position *i* transitioning to the next state of the Markov chain according to the transition probabilities of the class at position *i* (see Figure 1). A similar model was proposed by Burge and Karlin (1997) for gene finding; their hidden Markov model uses different transition and emission probabilities based on the GC content in the current window of the genome.

**Figure 1.**
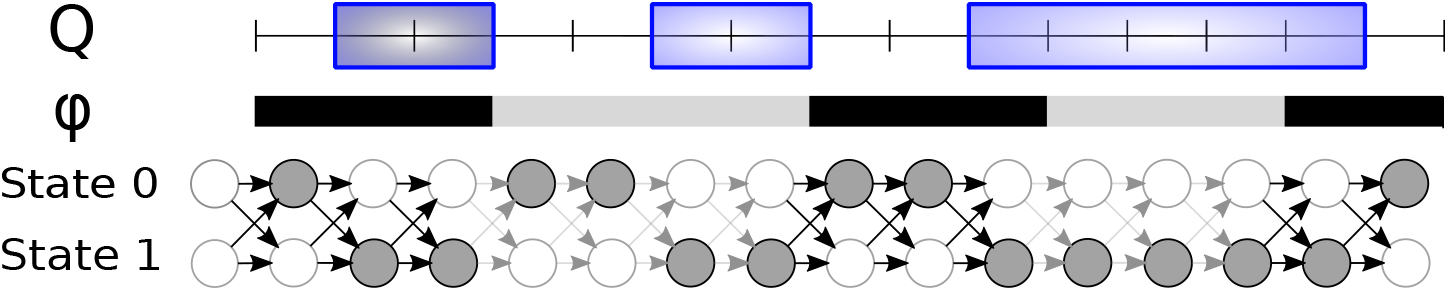
An example of a query annotation Q = {[1, 3), [5, 7), [9, 14)} and the corresponding sequence of states of the context-aware Markov chain that induces the annotation. Genome context ϕ is shown with black and gray colors corresponding to two distinct class labels. The same colors are also used on transition arrows between successive states of the Markov chain, as the transition probabilities depend on the genome context.

#### Definition 1.

*A* context-aware Markov chain *is a pair* (*ϕ*, **T**), *where ϕ is a genome context and* **T** : Λ *→* ℝ^2*×*2^ *is a mapping that provides a transition probability matrix for each context class. The context-aware Markov chain* (*ϕ*, **T**) *generates a sequence of states* (*s*_*−*1_, *s*_0_, …, **s**_*L−*1_) ∈ {0, 1}^*L*+1^ *with probability*

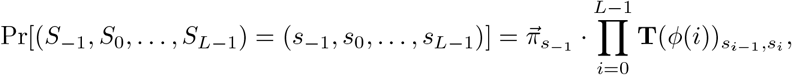

*where* **T**(*ϕ*(i))_*s,s′*_ *is the probability of transition from state s to state s^′^ in context class ϕ(i), and* 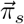 *is the probability of state s in the stationary distribution of the Markov chain with transition probabilities* **T**(*ϕ*(0)). *Specifically*,

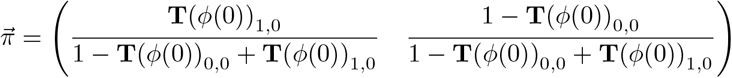

Note that we are indexing vectors and matrices starting from 0 in order to make the formulas more readable. The produced binary sequence of states (*s*_0_, …, *s*_*L−*1_) can be viewed as an annotation of a genome of size *L*. State *s*_*−*1_ is added to the start for notational convenience, as we will often refer to the state preceding the start of an interval. The distribution of the random vector of generated states (*S*_*−*1_, *S*_0_, …, *S*_*L−*1_) will be denoted as *C*(*ϕ*, **T**). We will use the same notation to denote the distribution of the induced annotation.

### 2.2 The context-aware Markov chain null model

The generative model above serves as a basis for our null model, which we call 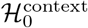. Given a context *ϕ* and a query annotation *Q* = (*Q*_0_, …, *Q*_*L−*1_), we first need to find the transition probabilities **T**_*Q*_ that maximize the probability of the context-aware Markov chain (*ϕ*, **T**_*Q*_) generating *Q*. This is achieved through the standard approach of training Markov chains by counting transition frequencies (Durbin et al., 1998). Namely, for each class, we count the number of times each possible state transition occurs in *Q*. Formally,

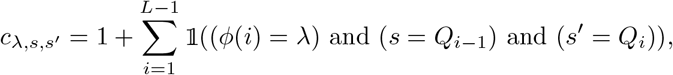

where 𝟙 is the indicator function which evaluates to 1 if the logical expression inside is true and 0 otherwise. A pseudocount 1 is added to avoid zero probabilities (Durbin et al., 1998). The transition matrix is then defined from these counts as

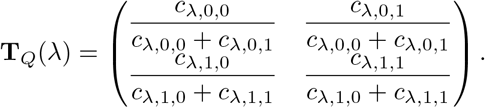

The mapping **T**_*Q*_ is computable in time 𝒪(|*Q*| + *c*) and space 𝒪(|Λ|), where c is the number of class boundaries. We can now formally define the context-aware Markov chain null hypothesis.

#### Definition 2.

*The context-aware Markov chain null hypothesis* 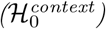 *for query annotation Q and context ϕ* : {0, …, *L−* 1} *→* Λ *is that the query annotation Q is generated by the context-aware Markov chain* (*ϕ*, **T**_*Q*_).

Note that under context *ϕ* with a single class, the context-aware Markov chain null hypothesis 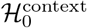 reduces to the Markov chain null hypothesis by Gafurov et al. (2022), with a small difference that the initial state distribution 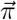 is set to the stationary distribution at position -1 instead of always starting in state 0 at position 0.

When there is just one context class,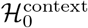 can be viewed as an approximation to the permutational null, i.e. shuffling the query intervals around in a random fashion (Gafurov et al., 2022). In the case of multiple classes, 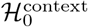 can be thought of as an approximation to shuffling the query intervals around separately within each class. However, 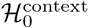 also transparently handles intervals spanning one or even multiple class boundaries.

### 2.3 Algorithm overview

We first state our main algorithmic result: fast computation of the mean and variance of *K*(*R, Q*) and *B*(*R, Q*) statistics under 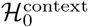. The mean and variance are then used to compute the p-values using the normal approximation.

#### Theorem 1.

*Let R and Q be two annotations and let ϕ be a genome context with* c *class boundaries. Let A be either the number of overlaps test statistic K or the number of shared bases test statistic B. It is possible to compute mean* E_*Q*′*∼C*_(*ϕ*,T_*Q*_)[*A*(*R, Q*^*′*^)] *and variance* Var_*Q*′*∼C*_(*ϕ*,T_*Q*_)[*A*(*R, Q*^*′*^)] *in space 𝒪*(|*R*| + *c*) *and in time*

- 𝒪 (|*R*| + |*Q*| + *c*) *when A is the overlap test statistics*,
- 𝒪 (|*Q*| + (|*R*| + *c*) log *t*) *when A is the shared bases test statistics; here, t is the length of the longest stretch of positions within a single reference interval with the same context class in R*.

Under the assumption that the test statistic is approximately normally distributed, this algorithm can be used to obtain the full probability mass function of its distribution under the null, i.e. values Pr_*Q*′*∼C*_(*ϕ*,T_*Q*_)[*A*(*R, Q*^*′*^) = *x*] for all values of *x*. Note that the previous algorithm for this problem MCDP only works with a single context, only works for the overlap statistic *K*, and runs in 𝒪(|*R*|^2^ + |*Q*|) time (Gafurov et al., 2022). However, it makes no assumption about normality.

In the rest of this section, we give an overview of our algorithm. Further details and proofs are provided in Section 2.4.To reuse parts of the algorithm for both *K* and *B*, we give a more general algorithm to compute the expectation and variance for a family of test statistics we call *separable* (not to be confused with *separable statistics* used by Medvedev (1977)).

#### Definition 3.

*Let R* = ([*b*_1_, *e*_1_), …, [*b*_|*R*|_, *e*_|*R*|_)) *be a reference annotation and let Q* = (*Q*_0_, …, *Q*_|*L−*1|_) *a query annotation expressed as a binary sequence. A test statistic A is called* separable *if there exists a function A*_*v*_ *accepting an arbitrary number of binary arguments such that we can express A as*

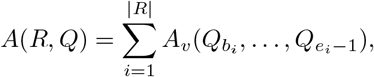

*We will refer to A*_*v*_ *as to the* interval function *of A and use it to define a separable test statistic*.

Note that both the number of overlaps and the number of shared bases are separable statistics with interval functions 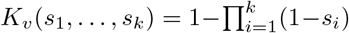 and 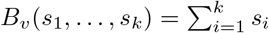, respectively.

Thanks to linearity of expectation, the expected value of any separable statistic can be computed for every reference interval separately and then summed together. The simplest case is the shared bases statistic B, which can be expressed as the sum of indicator variables for each base covered by *R*, and in the case of single-class null model, the expectation can be computed simply as the number of bases covered by *R* multiplied by the stationary probability of state 1 of the Markov chain. Context-aware models complicate the situation, as each base of the genome has its unique marginal distribution over states, depending on the sequence of class labels preceding it.

Computing variance is more complicated, as the values of the statistic in individual intervals of R are dependent, and therefore the overall variance is not a simple sum of individual variances. However, in a sequence of Markov chain states (*S*_0_, …, *S*_*L−*1_), states *S*_*i*_ and *S*_*j*_ are conditionally independent given *S*_*k*_ = *s* for *i ≤ k ≤ j*. Therefore, if random variable *X* is a function of *S*_*i*_, …, *S*_*k−*1_ and random variable *Y* is a function of *S*_*k*_, …, *S*_*j−*1_, then Var[*X* + *Y* | *S*_*k*_ = *s*] = Var[*X* | *S*_*k*_ = s] + Var[*Y* | *S*_*k*_ = *s*].

Our algorithm computes conditional variance in individual intervals of *R* conditioning on states at both interval boundaries, and then combines them using this formula. In order to remove conditioning on the boundary states, we use the law of total variance, which is summarized in Lemma 1.

The key data structure in our algorithm is a 𝒪(1)-sized vector called a *two-sided plumbus* defined below. It contains the quantities that we need to compute for every interval of *R*, conditioning on states at the interval boundaries. In the definition, function *v* expresses the contribution of a reference interval to the separable test statistics.

#### Definition 4.

*Let* (*ϕ*, **T**) *be a context-aware Markov chain with state sequence S*_*−*1_, *S*_0_, … *S*_|*L*|*−*1_, *let* [*i, j*) *be a subinterval of* [0, *L*), *and let v be a function on a binary sequence of length j − i. We define the* two-sided plumbus *for interval* [*i, j*) *as the collection of values*

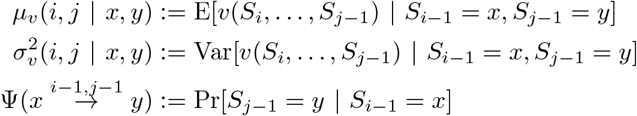

*for all combinations of* (*x, y*) *in* {0, 1}^2^.

The two-sided plumbuses computed for individual intervals of *R* and gaps between them are then combined to plumbuses for successively longer intervals, until we cover the whole chromosome and obtain the overall variance and expected value of the statistic of interest (see Figure 2).

**Figure 2.**
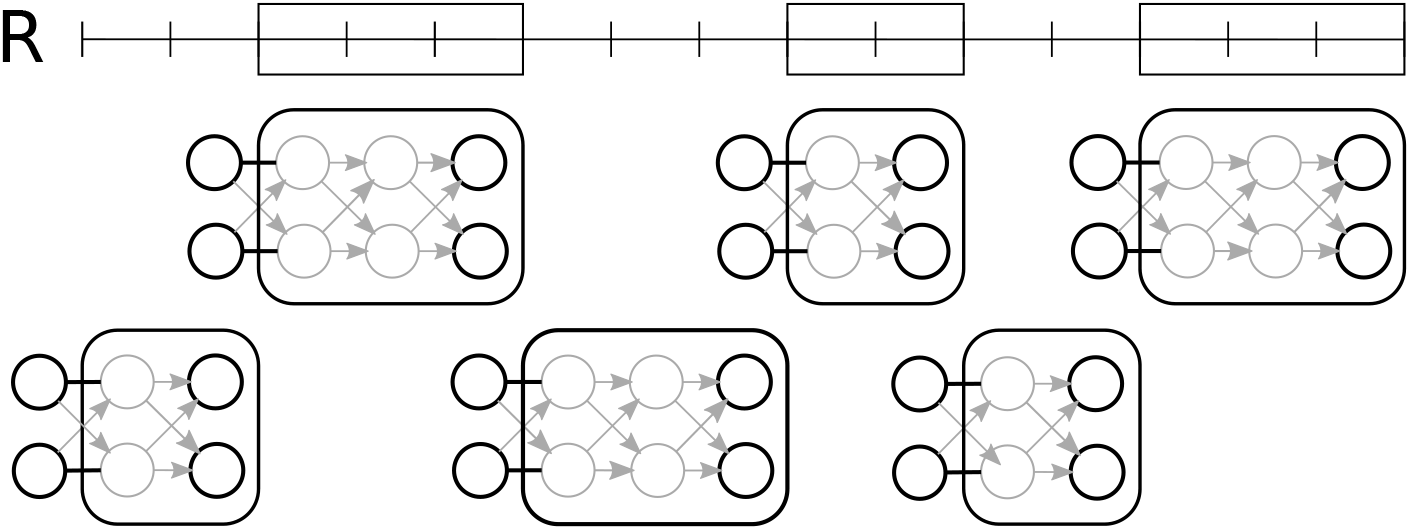
An example of a reference annotation *R* and the corresponding two-sided plumbuses. The plumbuses in the first row correspond to the reference intervals, and the plumbuses in the second row correspond to the gaps between the intervals. We highlight in black the boundary states on which we condition the values in each plumbus. Note that the conditional means *μ*_*v*_ and variances 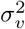 in the gap plumbuses are constant zeroes, since gaps do not contribute to the total test statistic.

In the algorithm, we compute Pr[*S*_*j*_ = *y* | S_*i*_ = *x*] and Pr[*S*_*i*_ = *S*_*i*+1_ = *· · ·* = S_*j*_ = 0] in constant time, provided that interval [i, j] is labeled by the same context class. This leads to a linear-time algorithm for *K*(*R, Q*) statistics. For *B*(*R, Q*) statistics, we split an interval of R into subintervals of size 1, compute plumbuses for them, and combine them in a similar manner, as we combine plumbuses in the overall algorithm. However, within a single context class, corresponding plumbus depends only on the interval length, and thus we can compute plumbuses for interval sizes which are powers of two and combine them to obtain a plumbus for any interval length within a single context in logarithmic time.

### 2.4 Algorithm details

In this section, we provide detailed description and proofs of individual parts of our algorithm in the form of a series of lemmas roughly corresponding to algorithm subroutines. The first lemma shows how to remove conditioning on a binary variable from expectation and variance using the law of total expectation and the law of total variance (also known as Adam’s law and Eve’s law). The lemma will be used repeatedly to compute and combine plumbuses.

#### Lemma 1.

*Let* A *be a random variable and B a binary random variable. If we are given values* Pr[*B* = *b*], E[*A* | *B* = *b*] *and* Var[*A* | *B* = *b*] *for b* ∈ {0, 1}, *we can compute* E[*A*] *and* Var[*A*] *in 𝒪*(1) *time*.

*Proof*. The expectation E[A] can be computed from the known values using the law of total expectation:

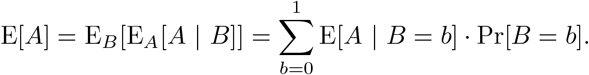

The variance equation can be obtained from the law of total variance:

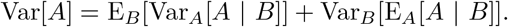

We now rewrite the first and second term separately as follows:

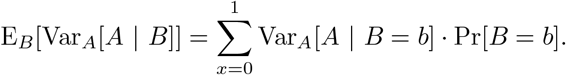

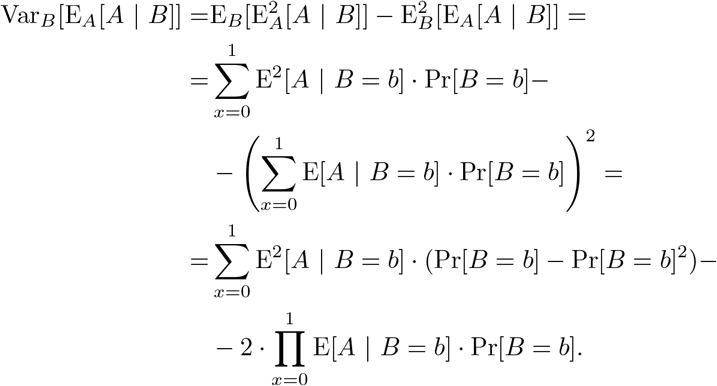

Clearly both terms can be computed in constant time from the provided values. □

Next, we introduce notation expressing conditional state probabilities in the context-aware Markov chain with state sequence *S*_*−*1_, *S*_0_, … *S*_|*L*|*−*1_. Let *x, y* ∈ {0, 1} and let *−*1 *≤ i ≤ j* < L. We define

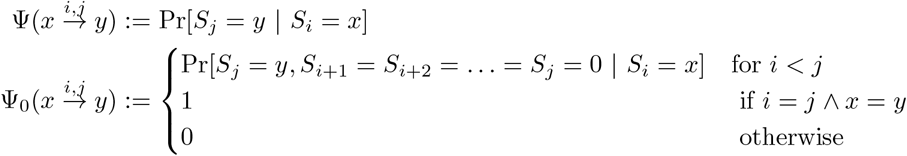

Value 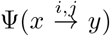 corresponds to the probability of transitioning from state *x* at position *i* to state *y* at position j. Function Ψ_0_ is similar to Ψ, but only state 0 is allowed from position *i* + 1 to *j*. Note that Ψ was already introduced in Definition 4, but is repeated here for completeness. The next lemma shows how to compute these quantities efficiently.

#### Lemma 2.

*Given a context-aware Markov chain* (*ϕ*, **T**), *values* 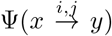 *and* 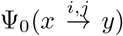 *can be computed in 𝒪*(c) *time and 𝒪*(1) *additional space, where* c *is the number of class boundaries between positions* i *and* j, *assuming that the class boundary positions are given*.

*Proof*. Both of those values can be represented using matrix multiplication:

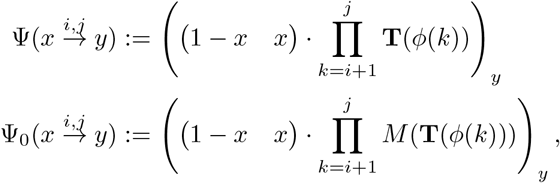

where *M* is a transformation that replaces the last column with zeros, i.e.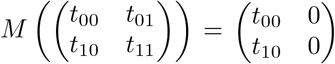. Note that (1 *− x x)* = (1 0) for *x* = 0 and (1 *− x x*) = (0 1) for *x* = 1.

Assume the class boundary positions are *t*_1_, …, *t*_*c*_. We also add *t*_0_ := i and *t*_*c*+1_ := *j* in order to unify the notation. Now, we can group the multiplied transition matrices into a product of exponentiations:

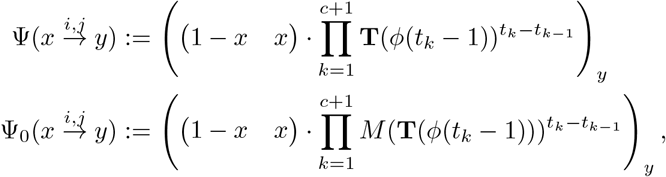

Matrices 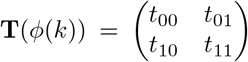 and 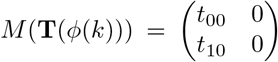 can be exponentiated to an arbitrary positive integer power a in time 𝒪 (1) using their diagionalizations (Gafurov et al., 2022):

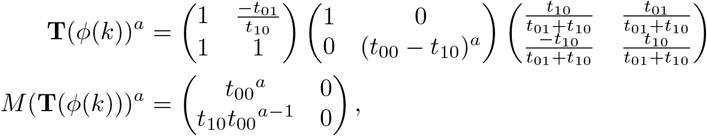

which concludes the proof of time and space complexity of the algorithm.

In addition to two-sided plumbuses from Definition 4, we will also compute one-sided plumbuses, which condition the mean and variance only on the state at the left side of the interval, as defined below.

#### Definition 5.

*Let* (*ϕ*, **T**) *be a context-aware Markov chain with state sequence S_−1_, S_0_, … S*_|*L*|−1_, *let* [*i, j*) *be a subinterval of* [0, *L*), *and let v be a function on a binary sequence of length j − i. We define the* left-sided plumbus *for* [*i, j*) *as the collection of values*

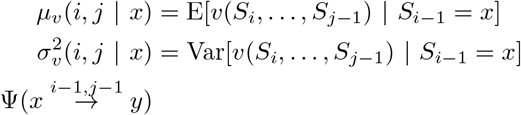

*for all combinations of* (*x, y*) *in* {0, 1}^2^.

Lemmas 3, 4 and 5 describe basic operations on one-sides and two-sides plumbuses that can be performed in constant time.

#### Lemma 3.

*(Conversion from two-sided to a left-sided plumbus) Given a two-sided plumbus, it is possible to compute the left-sided plumbus for the same function on the same interval in time 𝒪*(1).

*Proof*. Let the collection of values *μ*_*v*_ (*i, j* | *x, y*), 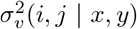 and 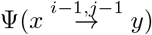 for all combinations of (*x, y*) in {0, 1}^2^ be a two-sided plumbus for function v on interval [*i, j*).

Transition probabilities in the corresponding left-sided plumbus are the same as in the two-sided plumbus, so we only need to compute values *μ*_*v*_(*i, j* | *x*) and 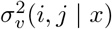 for all values of *x* in {0, 1}. We can compute them in constant time by applying Lemma 1 to marginalize variable y using values contained in the original two-sided plumbus. The input will be values 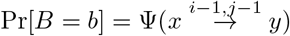, E[*A* | *B* = *b*] = *μ*_*v*_(*i, j* | *x, y*), 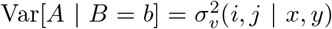 for *y* ∈ {0, 1}, and the result will be values E[A] = *μ*_*v*_(*i, j* | *x*) and 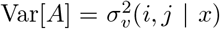. The computation is repeated for both values of x. □

#### Lemma 4.

*(Merging lemma for plumbuses) Given two two-sided plumbuses for functions f and g on intervals* [*i, j*) *and* [*j, k*), *respectively, it is possible to compute the two-sided plumbus for function h*(*S*_*i*_, …, *S*_*k−*1_) = *f*(*S*_*i*_, …, *S*_*j−*1_) + *g*(*S*_*j*_, …, *S*_*k−*1_) *on interval* [*i, k*) *in time 𝒪*(1).

*Proof*. Transition probabilities needed in the plumbus are computed by marginalizing over state *S*_*j*−1_:

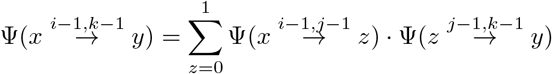

The expectations and variances are again computed using Lemma 1. Thanks to Markov property, we have

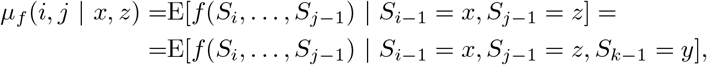

and similarly for *μ*_*g*_(*j, k* | *z, y*) and the corresponding variances. The expectation of function h conditioning on three states *x, z* and *y* is therefore

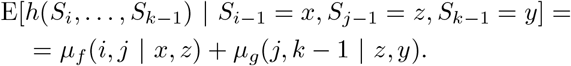

For variance we can also use addition as *f* and *g* are conditionally independent given *S*_*j−*1_ = *z*:

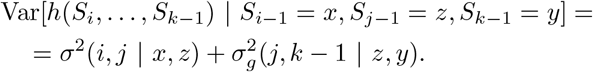

The final step is to use Lemma 1 to remove conditioning on value *z* and obtain *μ*_*h*_(*i, k* | *x, y*) and 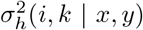. Probability Pr[*B* = *b*] needed by the lemma is computed as follows

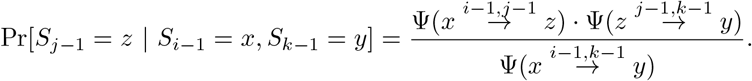

#### Lemma 5.

*(Folding lemma for plumbuses) Given a two-sided plumbus for function f on interval* [*i, j*) *and a left-sided plumbus for function g on interval* [*j, k*), *it is possible to compute the left-sided plumbus for function h*(*S*_*i*_, …, *S*_*k−*1_) = *f*(*S*_*i*_, …, *S*_*j−*1_) + *g*(*S*_*j*_, …, *S*_*k−*1_) *on interval* [*i, k*) *in time 𝒪*(1).

*Proof*. The computation proceeds similarly as in the previous lemma. We apply Lemma 1 to expected values *μ*_*f*_ (*i, j* | *x, z*)+*μ*_*g*_(*j, k* | *z*) and variances 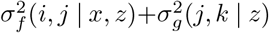 using probabilities 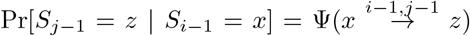 Value 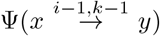 is computed as in the previous lemma.

The next two theorems show the details of efficient plumbus computation for the two statistics *K*(*R, Q*) and *B*(*R, Q*) considered in this paper.

#### Theorem 2.

*For given reference annotation R and context-aware Markov chain* (*ϕ*, **T**), *the two-sided plumbuses for interval function v of the number of overlaps statistic K on all intervals in R are computable in time 𝒪* (|*R*| + *c*) *and additional space 𝒪* (|*R*|).

*Proof*. Given interval [*b*_*j*_, *e*_*j*_) from R, random variable 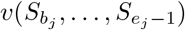 is an indicator variable, so its conditional mean and variance can be computed as *μ*_*v*_(*b*_*j*_, *e*_*j*_ | *x, y*) = 1 *− q*(*b*_*j*_, *e*_*j*_ | *x, y*) and 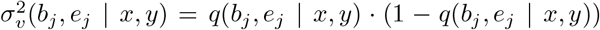, respectively, where 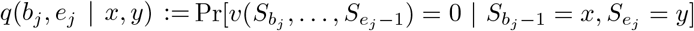. Value *q*(*b*_*j*_, *e*_*j*_ | *x, y*) can be computed as follows:

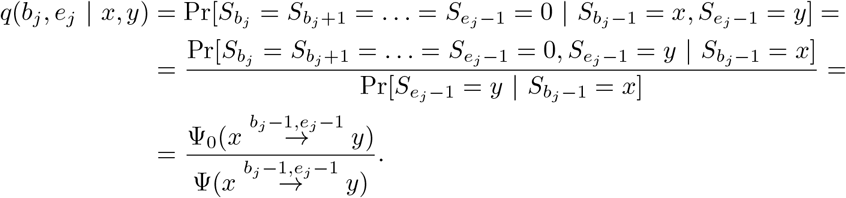

Assuming the class boundary positions are available sorted, and the reference intervals are sorted as well, it is then possible to compute values *q*(*b*_*j*_, *e*_*j*_ | *x, y*) for all combinations of the input arguments in time 𝒪(|*R*|+*c*) and space 𝒪(|*R*|), using fast computation of Ψ and Ψ_0_ values provided by Lemma 2. Since this computation already includes the computation of transition probabilities 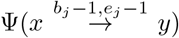 for each combination of values *x, y, j*, all the values required for the plumbuses are computable in the required time and space.

#### Theorem 3.

*For a given reference annotation R and context-aware Markov chain* (*ϕ*, **T**), *the two-sided plumbuses for interval function v of the number of shared bases statistic B on all intervals in R are computable in time 𝒪*((|*R*| + *c*) *·* log *t*) *and additional space 𝒪*(|*R*|), *where t is the length of the longest stretch of positions within a single reference interval with the same context class*.

*Proof*. Let *t*_1_, …, *t*_*d*_ be the class boundaries inside the *j*-th reference interval [*b*_*j*_, *e*_*j*_). Let us also define *t*_0_ := *b*_*j*_ and *t*_*d*+1_ := *e*_*j*_ in order to unify the computations. We will split interval [*b*_*j*_, *e*_*j*_) into d + 1 intervals [*t*_0_, *t*_1_), [*t*_1_, *t*_2_), …, [*t*_*d*_, *t*_*d*+1_) with a single context class within each individual interval (see Figure 3). Once we compute the two-sided plumbus for each interval [*t*_*i*_, *t*_*i*+1_), we can merge them using Lemma 4 into a two-sided plumbus on the whole *j*-th reference interval in time 𝒪(*d*).

**Figure 3.**
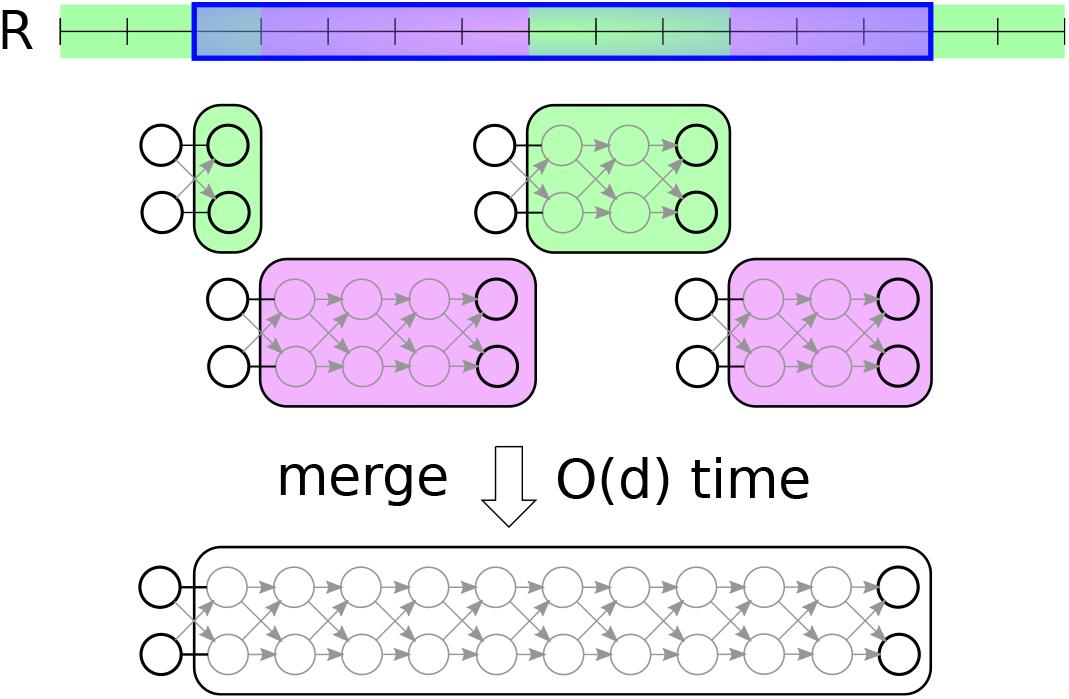
Example of decomposition of a single reference interval into single-class two-sided plumbuses. Note that the individual single-class two-sided plumbuses for the number of shared bases statistics B(·, ·) can be computed in time logarithmic to the length of their corresponding genomic interval and then merged into a single plumbus for the whole reference interval in time linear w.r.t. their count (see the proof of Theorem 3).

Let us now describe computation of the two-sided plumbus for a single-class interval [*t*_*i*_, *t*_*i*+1_) with class label λ. We can split the interval into *t*_*i*+1_ *− t*_*i*_ intervals of length 1 [*t*_*i*_, *t*_*i*_ + 1), [*t*_*i*_ + 1, *t*_*i*_ + 2), …, [*t*_*i*+1_ *−* 1, *t*_*i*+1_). The plumbuses on all those intervals are the same and can be computed in 𝒪(1) time:

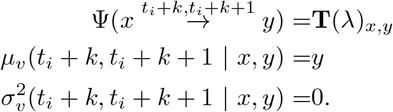

Similarly, two-sided plumbus for any subinterval of [*t*_*i*_, *t*_*i*+1_) of length *k* depends only on the length, not on its specific position. We can therefore use the merging operation from Lemma 4 to compute two sided plumbuses for intervals of lengths that are powers of two and to combine these to the two-sided plumbus for the whole interval [*t*_*i*_, *t*_*i*+1_) in time 𝒪 (log(*t*_*i*+1_ *− t*_*i*_)) and additional space 𝒪 (1).

Thus, we can compute two-sided plumbus on *j*-th reference interval in time 𝒪 ((*d* + 1) *·* log *t*) and additional space 𝒪 (1). Applying this to all reference intervals, we would obtain the desired time and space complexity of the algorithm.

Finally, the next theorem gives a general algorithm for computing mean and variance of any separable statistic. When we substitute results obtained for statistics *K* and *B* from Theorems 2 and 3, we obtain our main result from Theorem 1.

#### Theorem 4.

*For any separable statistic A, reference annotation R and context-aware Markov chain* (*ϕ*, **T**), *it is possible to compute mean* E_*Q*′*∼C*_(*ϕ*,T)[*A*(*R, Q*^*′*^)] *and variance* Var_*Q*′*∼C*_(*ϕ*,T)[*A*(*R, Q*^*′*^)] *in time* 𝒪 (|*R*| +*c*+*d*) *and space* 𝒪 (|*R*| +*c*+*e*), *where c is the number of class boundaries in context ϕ, and* 𝒪 (*d*) *and* 𝒪(e) *are the time and space complexity of computing two-sided plumbuses for the separable statistic A on each interval in the reference annotation R*.

*Proof*. We are provided two-side plumbuses for all reference intervals. To handle the gaps between these intervals, we will use zero interval function *z*(*s*_1_, …, *s*_*k*_) = 0. Using this function, we can obtain auxiliary two-sided plumbuses for all gaps in the reference annotation R including the potential gap between the start of the genome and the first reference interval. This works in total time 𝒪(|*R*| + *c*) and space 𝒪(|*R*|), since conditional means and variances are all equal to zero, and we only need to compute the transition probabilities using Lemma 2.

Then, we can convert the two-sided plumbus of the right-most reference interval into a left-sided plumbus in time 𝒪(1) (Lemma 3). Starting from that left-sided plumbus, we can then fold all interval and auxiliary plumbuses in time 𝒪(|*R*|) (Lemma 5). This would leave us with a left-sided plumbus for the test statistic A on interval [0, *e*_|*R*|_).

Finally, we can compute the unconditional mean E_*Q*′*∼C*_(*ϕ*,T)[*A*(*R, Q*^*′*^)] and variance Var_*Q*′*∼C*_(*ϕ*,T) [*A*(*R, Q*^*′*^)] of the test statistic *A* by applying Lemma 1 on the expected values *μ*_*A*_(0, *e*_|*R*|_ | *x*) and variances 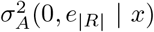 contained in this final left-sided plumbus. The probability of *S*_*−*1_ having value x is simply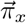 where 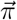 is the initial state distribution of context-aware Markov chain (*ϕ*, **T**).

Adding up the total time and space complexity with the time and space requirements to compute the plumbuses for the reference intervals, we obtain the time and space complexities as given in the theorem.

### 2.5 Multiple chromosomes

Both our model and our algorithm can be extended to genomes with multiple chromosomes in a straightforward way. We assume that the query annotation is generated independently for each chromosome. The training of the context-aware Markov chain is accomplished simply by counting transition frequencies on all chromosomes. The test statistic for the whole genome is defined as the sum of test statistic values for the individual chromosomes. This, in turn, allows us to compute the mean and variance of the total statistic by summing the means and variances, respectively, for the individual chromosomes. Note that this simple computation works for the variance thanks to the chromosome independence assumption. Therefore, the time and space complexity of our algorithm remains the same for the case of multiple chromosomes.

## 3 Experiments

### The normal distribution yields an accurate p-value approximation

Our MCDP2 algorithm computes the exact expectation and variance of the null distribution and uses them to approximate the null distribution by the normal distribution. Here, we first compare the accuracy of this approximation for the *K*(*R, Q*) statistic with the exact distribution computed by the previous MCDP algorithm (Gafurov et al., 2022). The comparison was performed on synthetic data sets with genome length *L* = 10^8^ bp, query annotations with 50 000 randomly generated intervals of length 500 bp each and reference annotations with up to 20 000 intervals of length 500 bp each. To understand the influence of the number of reference intervals on the accuracy, we vary |R| from 200 to 20000.

Figure 4 shows that the exact PMF in general agrees well with the normal approximation. The approximation approach allows to estimate even very low p-values accurately with the growing number of reference intervals. However, for |*R*| = 200 the differences in the extreme tail of the distribution lead to overly conservative p-values. Therefore, for small values of |*R*| we recommend the use of the exact MCDP algorithm, which is not time-prohibitive. The new MCDP2 tool includes a reimplementation of the exact computation of the PMF and its extension to multiple context classes, and we use it in these experiments under the label MCDP*.

**Figure 4.**
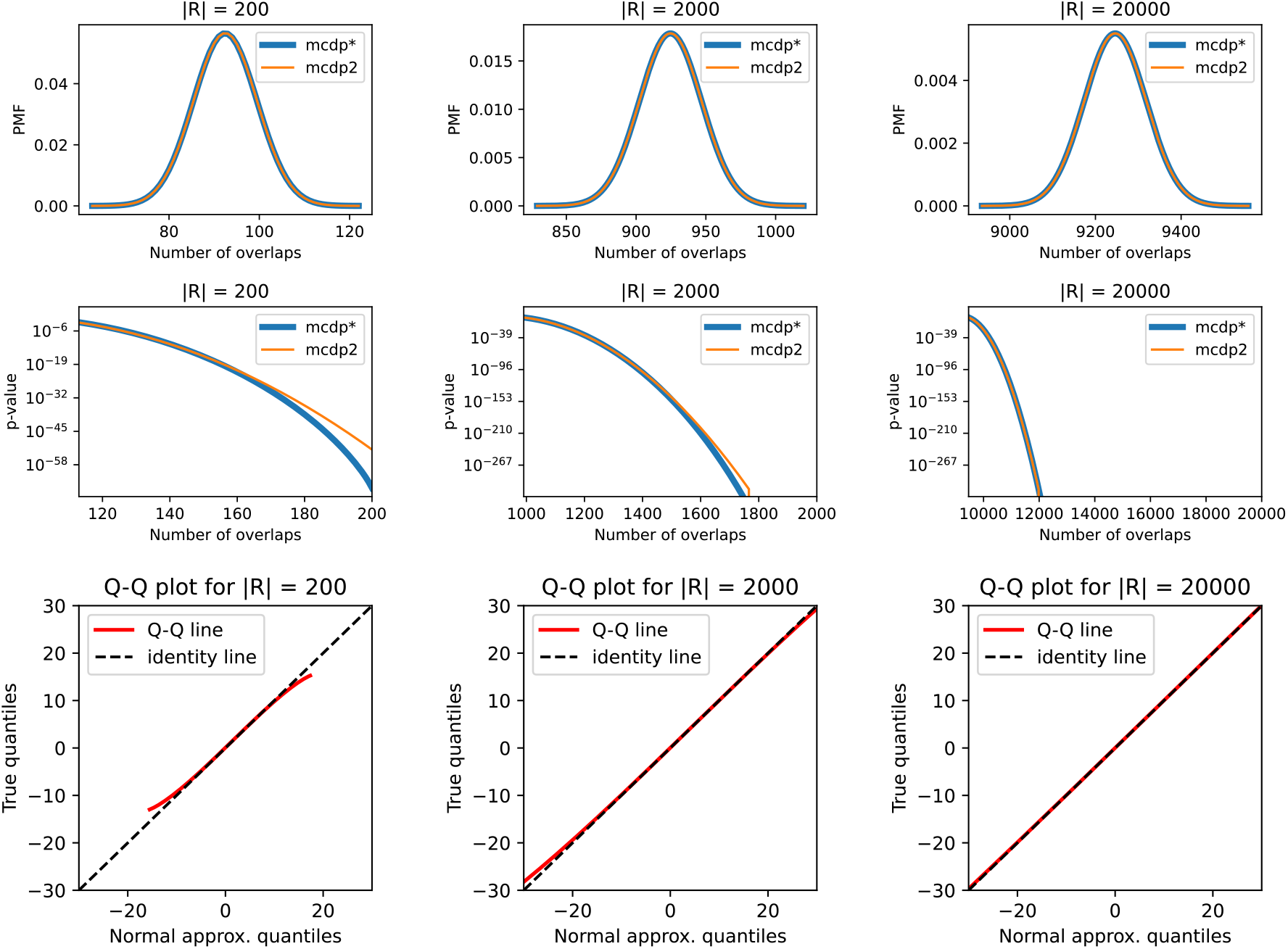
The comparison of the exact probability mass function (PMF) for the number of overlaps *K* statistic (MCDP^*^) with its normal approximation (MCDP2) on synthetic data sets. Each column represents a different number of reference intervals (|*R*| ∈ {200, 2000, 20000}). The top row compares the central part of the two distributions. The middle row shows differences for the extreme tail of the distributions. The curves represent the exact and approximated p-value for different values of the statistic, starting from the position with Z-score +3. Finally, the bottom row contains the quantile-quantile plots (Q-Q plots) between the two distributions.

We also tested the accuracy of our normal approximation for the number of shared bases *B* statistic in comparison with the full PMF computed by an exact algorithm. To address the fact that the original MCDP algorithm does not work for this statistic, we split each reference interval into a sequence of adjacent intervals of size one, producing a modified reference annotation *R*^*′*^. For example, reference annotation *R* = {[2, 5), [7, 9)} would yield *R*^*′*^ = {[2, 3), [3, 4), [4, 5), [7, 8), [8, 9)}. It is easy to see that *B*(*R, Q*) = *K*(*R*^*′*^, *Q*), and we can therefore compute the exact PMF for *B*(*R, Q*) by computing the PMF for the *K*(*R*^*′*^, *Q*) statistic by the MCDP* algorithm, which was adjusted to allow adjacent intervals in the annotation (we otherwise assume that intervals are separated by at least one base). A major limitation is that the running time of MCDP is quadratic with respect to the number of reference intervals |*R*|. For this test, this turns into a quadratic dependence on the total number of *bases* in reference intervals, which we denote B(*R*). This limits our experiments to reference annotations with up to tens of thousands of bases.

The comparison was performed on synthetic data sets similar to those used for the number of overlaps *K*, scaled down to limit the total number of bases in the reference. The synthetic data sets were generated with genome length *L* = 10^5^ bp, query annotations with 500 randomly generated intervals of length 50 bp each. The reference annotations were generated with up to 400 intervals of length 50 bp each. To understand the influence of the total number of reference bases *B*(*R*) on the accuracy, we vary *B*(*R*) from 200 to 20 000, i.e. from 4 to 400 reference intervals.

Figure 5 shows the rapid convergence of the exact PMF to its normal approximation. The case with four reference intervals (the first columns of the figure) shows prominent peaks in the exact PMF, corresponding to probabilities of hitting zero, one, two or more complete reference intervals. The periodicity of the peaks is equal to the length of the reference intervals (50 bp). Nonetheless, forty intervals (the second column) are in this case enough for the PMF to be very close to normal around the center of the distribution. However, even with 400 intervals the PMFs still differ significantly in the extreme tails. We expect that approximation of even these extreme values is better for larger reference annotations which could not be tested due to computational costs of the full PMF computation. In practice we usually work with much larger annotations than the 400 intervals tested in this comparison.

**Figure 5.**
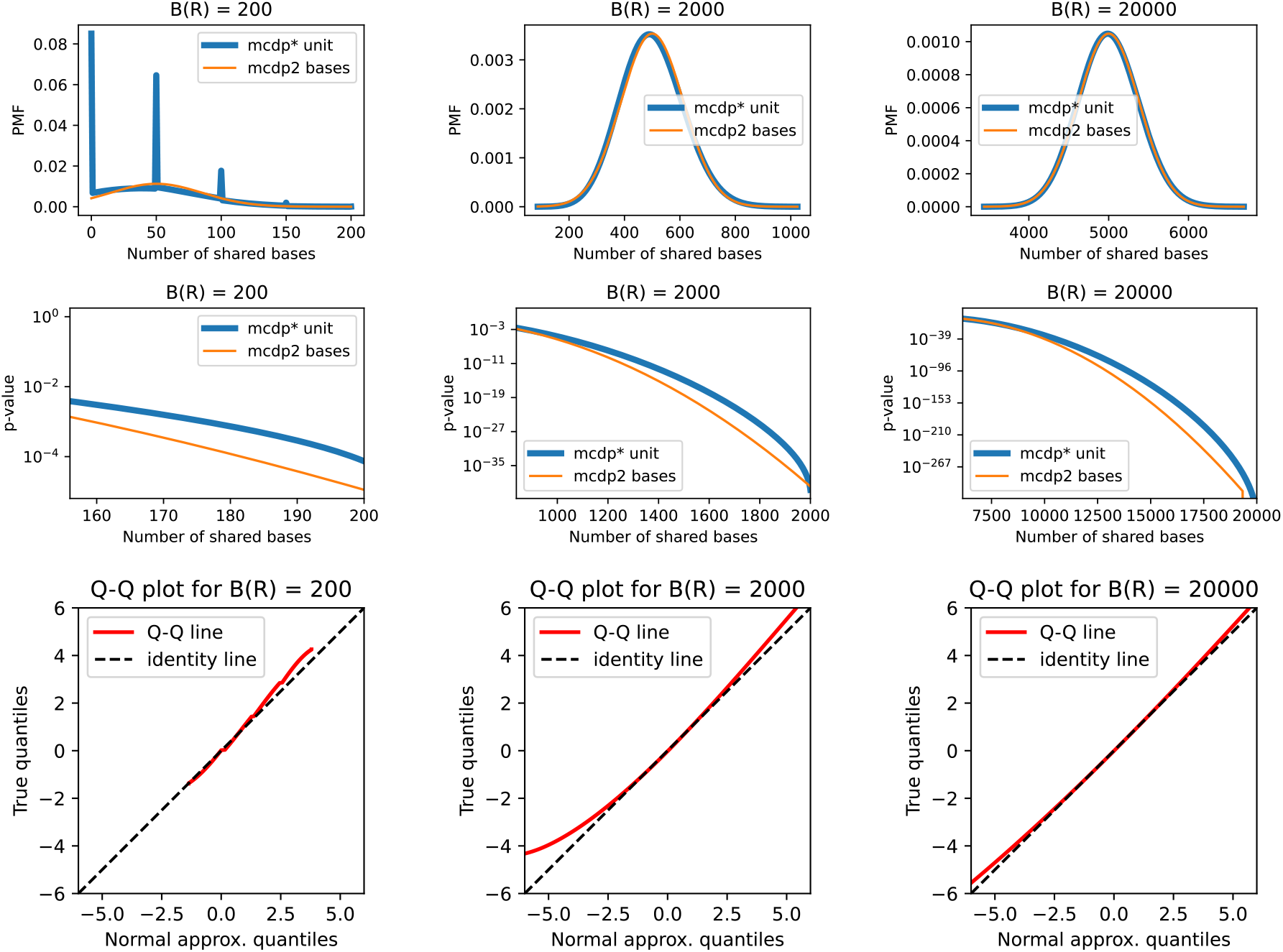
The comparison of the exact probability mass function (PMF) for the number of shared bases statistic B with its normal approximation on synthetic data sets. Each column represents a different total number of bases in reference intervals (*B*(*R*) ∈ {200, 500, 2000, 20000}). The top row compares the central part of the exact PMF (labeled “mcdp^*\**^ unit”) with the normal approximation computed using our new MCDP2 algorithm (labeled “mcdp2 bases”). The middle row shows p-values for the extreme tail of the distribution, starting from the position with Z-score +3. Finally, the bottom row contains the quantile-quantile plots.

### MCDP2 is fast and memory efficient

The speed of our algorithm enables us to apply our tools to large-scale comparisons, such as the data from a recent study of ENCODE epigenetic modification enrichment for different repeat types in the human genome (Gershman et al., 2022), employing the Telomere-to-Telomere (T2T) human genome assembly (Nurk et al., 2022). We use a context with two classes, one corresponding to all repeats and one to the rest of the genome, leading to over 4 million class boundaries. We use one of the 10 repeat types as the reference and one of the 45 available combinations of an epigenetic modification and a cell line as the query. Using 24 CPU threads, MCDP2 computed p-values for all 450 pairs in approx. 2 hours (wall clock) for the number of overlaps and approx. 3 hours for the number of shared bases, using at most 4.2 Gb (2.3 Gb) memory per comparison for overlaps (shared bases) statistic (see Table 1).

**Table 1:** Data set sizes and average running times for comparison of repeat types (*R*) with ENCODE epigenetic modifications (*Q*), using all repeats as context. Averages are computed across 45 different query annotations, each representing a specific epigenetic modification in a specific cell line. Note that the running time grows with |*R*| + *c*, and in this experiment, *c* is large even for inputs with small |*R*|.

| Repeat type | $\vert R\vert $ | time (s) | |
| --- | --- | --- | --- |
| | | $K(R, Q)$ | $B(R, Q)$ |
| RNA | 11 139 | 220 | 160 |
| Other | 8 835 | 220 | 157 |
| Unknown | 11 229 | 226 | 159 |
| Satellite | 47 041 | 229 | 202 |
| Low-complexity | 102 521 | 244 | 223 |
| DNA | 505 896 | 343 | 588 |
| LTR | 660 823 | 387 | 769 |
| Simple | 708 565 | 398 | 637 |
| LINE | 1 440 792 | 578 | 1 477 |
| SINE | 1 672 984 | 640 | 1 637 |

We compare the running time of MCDP2 to the quadratic-time MCDP* algorithm on the synthetic data sets used for Figure 4, with 20 pairs of *R* and *Q* generated for each setting (Figure 6). Note that MCDP* is faster than the original MCDP implementation (Gafurov et al., 2022), thanks to more extensive use of numpy library and reimplementation of part of the algorithm in C++. For the overlaps statistic, the MCDP* needs more than 1 000 seconds for 20 000 reference intervals, while our new approach MCDP2 only takes approximately 8 seconds on the same inputs. Computation for the number of shared bases is slightly slower (23 seconds for |*R*| = 20 000), which is consistent with its quasi-linear time complexity (in contrast to purely linear for the number of overlaps).

**Figure 6.**
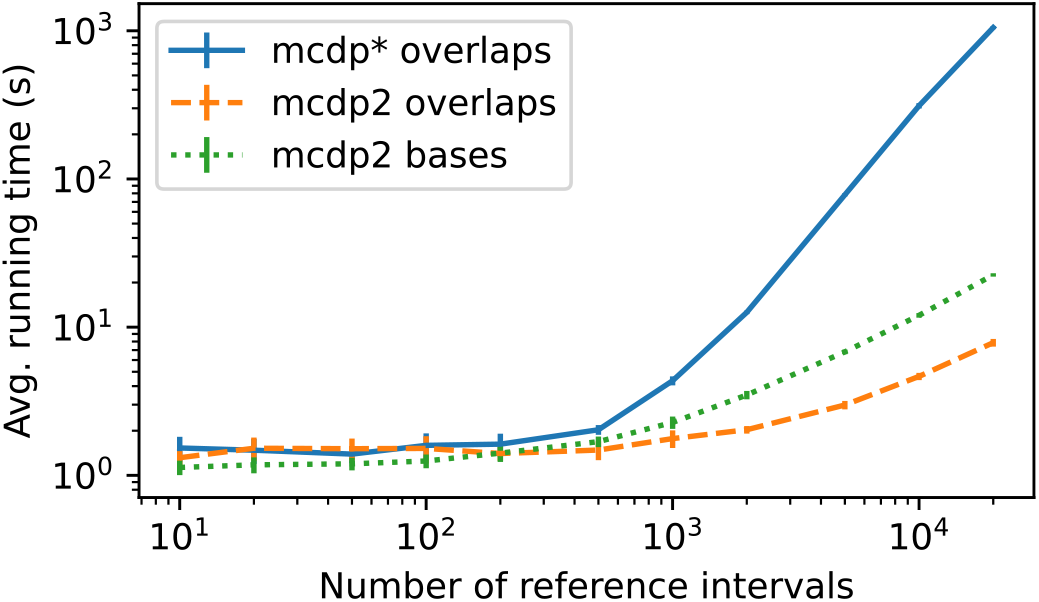
Average running time on synthetic data for overlaps statistics (MCDP2, MCFP*) and shared bases statistics (MCDP2). The vertical bars represent the standard deviation over 20 samples. Both axes are in log scale. The calculations were performed on a single thread on Intel(R) Xeon(R) Gold 6248R CPU.

### Genome contexts enable more detailed analysis of colocalization of copy number loss with different gene groups

To illustrate the power of our context-aware null model, we have reanalyzed the colocalization of exons of different gene groups with copy number loss regions, originally performed by Zarrei et al. (2015). Figure 7 shows the Z-scores for both *K* and *B* test statistics and for three types of contexts. The first context function only uses a single class. The second context function creates two classes by masking regions that are assembly gaps; this is motivated by the fact that both copy number losses and exons are annotated exclusively outside the gaps and, therefore, may appear colocated even if they were independent (previously also studied by Domanska et al. (2018)). The third context function uses six classes: one for gaps and the other five for discretization of the GC content in 1 kbp windows; this is motivated by the fact that GC content is known to be a significant confounding factor in many genomic analyses (Burge and Karlin, 1997; Gelfman and Ast, 2013).

**Figure 7.**
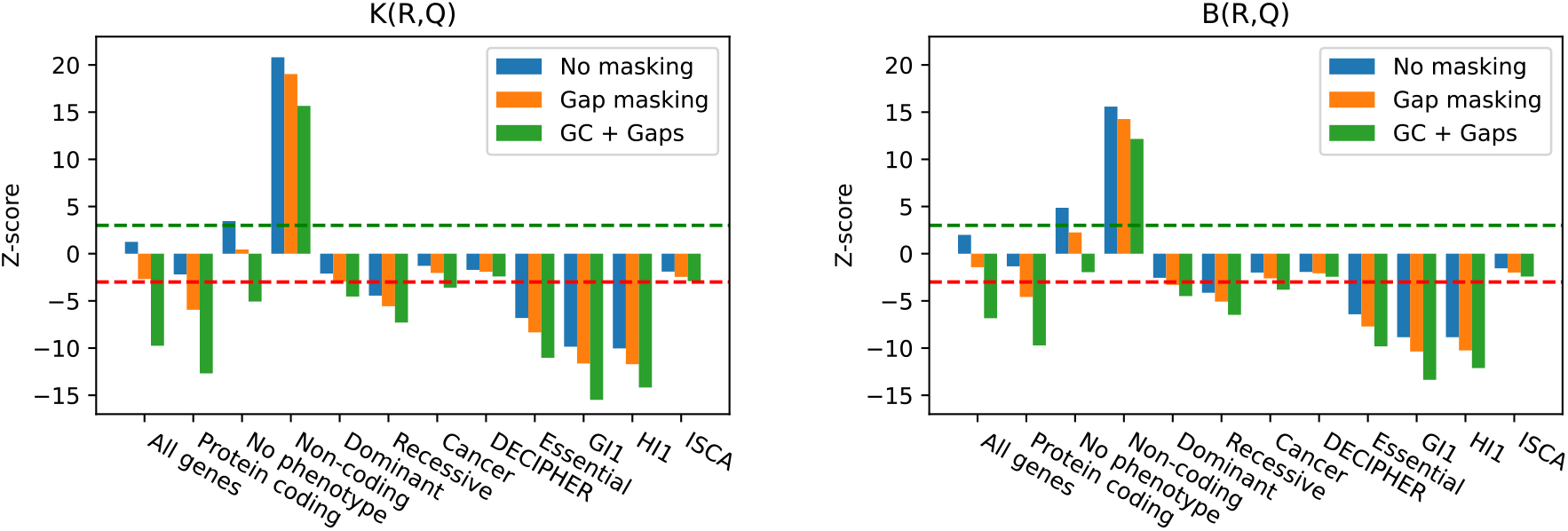
Z-scores for colocalization of exons of various gene groups (*R*, x-axis) with copy number losses (*Q*) under three different null models: single-class context, gap-aware context, and GC– aware context. Left: overlap statistics *K*; Right: shared bases statistics *B*. The green and red dashed lines stand for Z-score +3 and -3 respectively, corresponding to p-value 0.00135 for enrichment/depletion.

Figure 7 illustrates the importance of having a class in the context dedicated to gaps. In one jarring scenario, the set of all exons is enriched for overlaps with copy number losses (also observed by Zarrei et al.), but after accounting for gaps, the exons become depleted. More generally, across all studied gene groups, the Z-score decreases when the gaps are taken into account. This is expected as neither exons nor copy number losses occur in gaps, and thus ignoring gaps in the analysis may create spurious enrichments or lower the degree of observed depletion compared with analysis that takes gaps into account.

The GC-aware context also proves crucial for an accurate analysis. For example, the depletion of all exons for overlaps with copy number losses becomes much more pronounced in the GC-aware context. We provide a more detailed analysis of this result below. In another striking example, genes with no known phenotype appear enriched for overlap with losses (also in agreement with Zarrei et al.) when using the gap-aware context, but enrichment turns into slight depletion after taking GC content into account.

Other observations from Figure 7 are generally consistent with biological expectations. Protein coding genes are only slightly depleted in the single-class context but become significantly depleted when using gap-aware or GC-aware contexts. This depletion is consistent with the expectation that protein-coding exons are mostly evolutionarily conserved. Interestingly, the set of non-coding genes is strongly enriched for copy number losses under all three context functions, and the enrichment was also observed by Zarrei et al.

### Details of overlaps between all exons and copy number losses

In this section we show the details of the distribution of all exons (reference annotation *R*) and copy number losses (query annotation *Q*) and their overlaps in different genomic contexts. As we already discussed, the MCDP analysis without contexts as well as the original analysis by Zarrei et al. (2015) indicates enrichment, adding gaps turns the MCDP2 result into a slight depletion and adding five GC content class labels plus gaps leads to an even more pronounced depletion (Figure 7, label “All genes”). Detailed statistics are shown in Table 2. The five GC labels were chosen so that each corresponds to roughly the same amount of genomic sequence. The number of exon bases is similar in the first four GC classes, but it is much higher in the class with the highest GC. Copy number losses are also most frequent in high GC regions, but the difference is less pronounced.

**Table 2:**
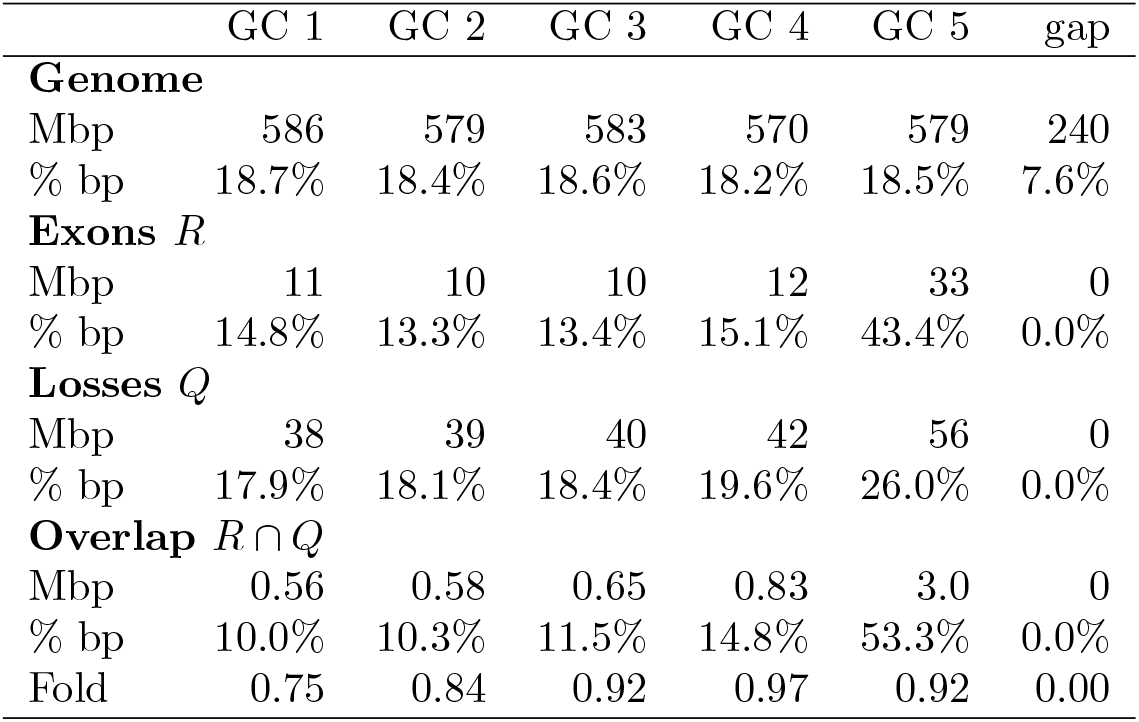
Detailed statistics for reference annotation *R* consisting of exons of all genes and query annotation *Q* consisting of copy number losses in the model with six context classes. For each context class, we show the number of bases from the whole genome, *R, Q* and *R* ∩ *Q* and what percentage of these bases is in each context class. Finally we also the simple fold change.

|  | GC 1 | GC 2 | GC 3 | GC 4 | GC 5 | gap |
| --- | --- | --- | --- | --- | --- | --- |
| **Genome** |  |  |  |  |  |  |
| Mbp | 586 | 579 | 583 | 570 | 579 | 240 |
| % bp | 18.7% | 18.4% | 18.6% | 18.2% | 18.5% | 7.6% |
| **Exons $R$** |  |  |  |  |  |  |
| Mbp | 11 | 10 | 10 | 12 | 33 | 0 |
| % bp | 14.8% | 13.3% | 13.4% | 15.1% | 43.4% | 0.0% |
| **Losses $Q$** |  |  |  |  |  |  |
| Mbp | 38 | 39 | 40 | 42 | 56 | 0 |
| % bp | 17.9% | 18.1% | 18.4% | 19.6% | 26.0% | 0.0% |
| **Overlap $R \cap Q$** |  |  |  |  |  |  |
| Mbp | 0.56 | 0.58 | 0.65 | 0.83 | 3.0 | 0 |
| % bp | 10.0% | 10.3% | 11.5% | 14.8% | 53.3% | 0.0% |
| Fold | 0.75 | 0.84 | 0.92 | 0.97 | 0.92 | 0.00 |

For each context class label, we calculated a simple fold change measure defined as the ratio between the observed and expected number of bases shared between *R* and *Q* under a simplified model with independent bases. For a context class with *t* bases in total, *r* reference bases and q query bases, the expected number of shared bases is 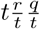. Our results show that even after taking into account different density of losses and exons in different GC contexts, the overlaps are depleted in all GC contexts, with the strongest 0.75-fold depletion in the lowest GC level and much weaker 0.92-0.97-fold depletion in the highest three GC content levels.

### Differential analysis of non-telomeric and telomeric TAR elements

Completion of the previously inaccessible parts of the human genome (Nurk et al., 2022) has allowed Gershman et al. (2022) to study telomere-associated repeats (TARs) and their colocalization with epigenetic modifications. While TARs located in subtelomeric regions are presumed to be important for telomere length regulation, TAR copies have also been dispersed to other parts of the genome. Differences between these two groups may further clarify mechanisms and functions of TARs in subtelomeric regions. While Gershman et al. observe differences in enrichment of some epigenetic marks, they do not assign statistical significance to their findings.

We adapted our context-aware Markov chain model to perform such differential analysis of enrichment between two annotations. In general, consider two references *R*_1_ ⊆ *R*_2_. In our case, *R*_1_ are non-telomeric TARs, *R*_2_ are all TARs, and *Q* are regions with a particular epigenetic mark. One could compare the enrichment p-value of *Q* in *R*_1_ with the enrichment p-value of *Q* in *R*_2_; however, this is not statistically sound (Goodman, 2008; Sullivan and Feinn, 2012). Instead, we create a context *ϕ*_rel_ with two class labels {outside, inside}, where positions covered by *R*_2_ are labeled “inside” and all other positions are labeled “outside.” We then use a test statistic to measure the significance of enrichment of *Q* in reference *R*_1_ with context *ϕ*_rel_. This context ensures that within *R*_1_ the null model uses the parameters estimated from intervals of *Q* that overlap *R*_2_, thus comparing colocalization of *Q* in *R*_1_ relative to colocalization of *Q* in the whole *R*_2_. Note that the query intervals can occur also outside of *R*_2_, and their properties are summarized in the parameters of the Markov chain for the “outside” class. These outside areas then influence the distribution of the test statistic under the null only by influencing the initial state distribution at the start of each interval of *R*_1_.

Figure 8 shows the result of this analysis applied to colocalization of epigenetic marks in non-telomeric TARs compared to all TARs. Similarly to Gershman et al. (2022), we observe relative enrichment of activating marks *H3K27ac* and *H3K4me3* in non-telomeric TARs using both *K* and *B* statistics. We can also see enrichment of *CTCF*, which is significant only under the shared bases statistic, perhaps due to the small number of intervals in *R*_1_. Gershman et al. were not able to observe relative enrichment for *CTCF* on non-telomeric TARs, although they do observe that *CTCF* is strongly enriched in both TAR classes compared to the background. This highlights usefulness of our context model in scenarios requiring relative analysis of two reference annotations.

**Figure 8.**
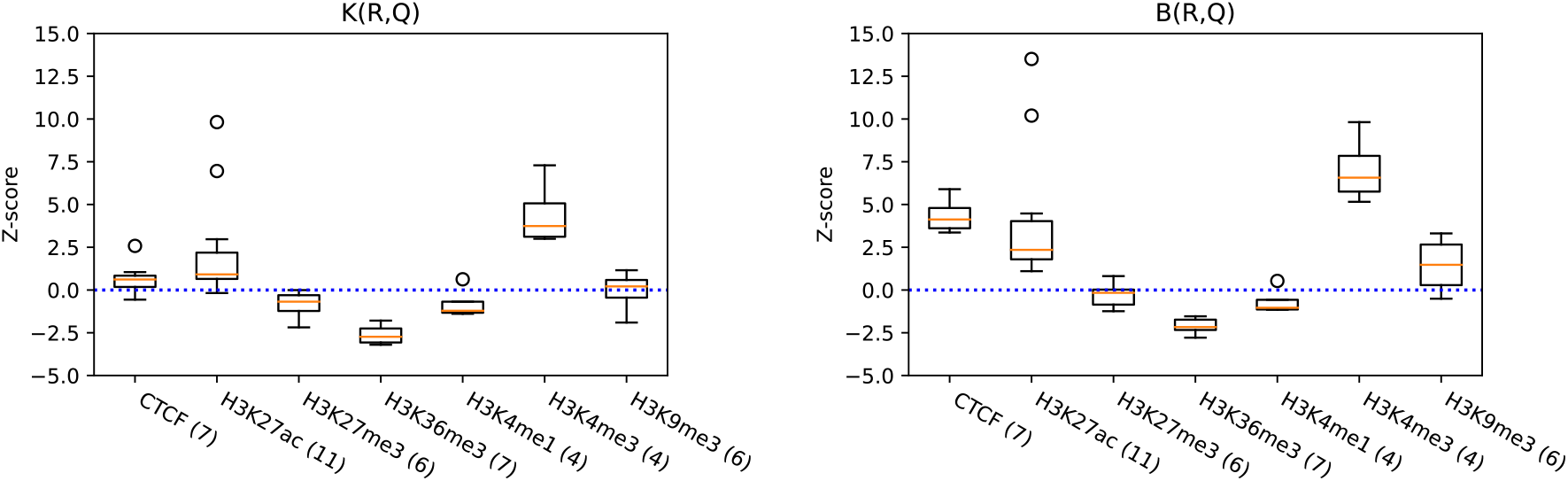
Relative enrichment of telomere-associated repeats (TARs) located further than 20 kbp from chromosome ends (*R*_1_) with epigenetic modifications (*Q*) in comparison to all TARs (*R*_2_), using both the number of overlaps statistic *K* (left) and the number of shared bases statistics *B* (right). The numbers in the parentheses denote the number of cell lines available for each modification.

## 4 Discussion

We have introduced a novel model for annotation colocalization analysis, which uses genomic contexts to capture confounding factors that may lead to false significance results. Taking advantage of the Markovian properties of our model, we have provided a general framework to compute the exact mean and variance of a broad class of colocalization test statistics (which we named *separable*). Using this framework, we were able to obtain linear and quasi-linear algorithms to compute the Z-scores for the number of overlaps and the number of shared bases. We have then proposed to convert the exact Z-score to approximate the p-values using the normal distribution.

Our algorithm computes a Z-score in 𝒪(|*Q*| + |*R*| +*c*) time for the overlap number statistic and in 𝒪(|*Q*| + (|*R*| + *c*) log *t*) time for the shared bases statistic, where |*Q*| and |*R*| are the number of intervals in the query and reference, respectively, *c* is the number of context class switches along the genome, and *t* is an upper bound on the reference interval length. This is in contrast to the previous best algorithm, which did not account for genome contexts and took 𝒪(|*R*|^2^ + |*Q*|) time to compute the probability mass function of the p-values.

In our experiments, we have demonstrated that our algorithm is sufficiently fast to allow large-scale studies comparing many pairs of annotations with large reference sets and frequent context class boundaries. We have reanalyzed data sets from two large-scale studies (Gershman et al., 2022; Zarrei et al., 2015), and thanks to our new context-aware model, we were able to further illuminate the nature of colocalizations discovered in these works, in some cases reversing previously published findings.

We have experimentally shown that the normal approximation of the distribution of the number of overlaps under 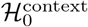 yields accurate p-values, and the approximation gets tighter with increasing number of reference annotation intervals. This behaviour intuitively follows from the representation of a separable statistic as a sum of contributions for individual reference intervals.

If those contributions were independent, their sum would converge to a normal distribution with a growing number of reference intervals under the classical central limit theorem. Though the contributions of individual intervals are dependent in our case, the fact that the dependencies stem from a Markov chain makes it possible that the sum converges under some extensions of the central limit theorem. In future, we hope to characterize sufficient conditions for such convergence. Additionally, we would like to explore the possibility of providing lower and upper bounds on the precision of the p-value estimation, possibly by applying the Stein’s method (Ross, 2011).

On a more practical side, in our future research, we would like to explore the possibility of using quantitative contexts, with numeric values such as GC content, epigenetic mark density, sequence conservation etc. Some work in this direction has already been done, particularly in HyperBrowser (Sandve et al., 2010). In MCDP2 this could be achieved for example by parameterizing the weights of the underlying Markov chains with the context value at each position. The challenge would be to keep the running time efficient for large genomes.

Another challenge is to provide statistical significance for statistics comparing colocalization of query *Q* with respect to two different references *R*_1_ and *R*_2_, such as *B*(*R*_1_, *Q*)/*B*(*R*_2_, *Q*). This may in some situations be preferable to our approach of comparing such colocalization through contexts, which we used for the analysis of TAR elements.

## Acknowledgments

Earlier versions of this article were published at bioRxiv and in the proceedings of the RECOMB 2024 conference.

